# The smORF-containing gene *mille-pattes* is required for moulting and *Trypanosoma cruzi* metacyclogenesis in the Chagas disease vector *Rhodnius prolixus*

**DOI:** 10.1101/2020.10.06.324897

**Authors:** Carina Azevedo Oliveira Silva, Sandy da Silveira Alves, Bruno da-Costa-Rodrigues, Jonatha Anderson Fraga Egidio, Lupis Ribeiro, Carlos Logullo, Flavia Borges Mury, José Roberto da Silva, José Luciano Nepomuceno-Silva, Rodrigo Nunes-da-Fonseca

## Abstract

Chagas disease is estimated to affect 8 million people worldwide and is responsible for circa 10,000 deaths in Latin America every year. Vector control has been developed as the main strategy to control disease spreading. The identification of new targets is essential, and genes containing small open reading frames (smORFs - < 100 amino acids) are present as hundreds of putative new candidate targets in insect genomes. Here, we show that the prototypic smORF containing gene *mille-pattes/polished-rice/tarsalless* (*mlpt/pri/tal*) is essential for post-embryonic development of the kissing bug *Rhodnius prolixus* and for the metacyclogenesis of the *Trypanosoma cruzi* parasite during nymphal stages. Injection of double-stranded RNA against *mlpt* (Rp-ds*mlpt*) during nymphal stages leads to a plethora of phenotypes which impairs post-embryonic development. First, nymphs of fourth or fifth stage injected with Rp-ds*mlpt* do not moult. Second, vector digestive physiology is largely modified, and haemozoin is significantly increased in the posterior midgut of Rp-ds*mlpt* nymphs. Third, Rp-*mlpt* knockdown inhibits the metacyclogenesis of *Trypanosoma cruzi*, the etiologic agent of Chagas disease. Thus, our study provides the first evidence of a smORF-containing gene regulating vector physiology, parasitic cycle and disease transmission.

## Introduction

Chagas disease was originally described in the American continent in the first decade of the 20th century as an infection caused by the flagellated parasite *Trypanosoma cruzi* and transmitted by contact with excrements from hematophagous triatomine bugs [1]. This anthropozoonosis is endemic to the American continent and recent estimates indicate that 6-7 million people are currently infected with *T. cruzi* worldwide, and 75 million individuals in Latin America are at risk of being infected [2]. American trypanosomiasis evolves first as a difficult to diagnose acute disease with mild and unspecific symptoms, which is followed by a chronic disease, mainly characterised by cardiomyopathy and arrhythmias, or by impairment of gastrointestinal function [3]. Despite several decades of research efforts directed to understand and combat Chagas disease, the control of vectorial transmission is still one of the main foci of public health approaches on this topic, and many questions remain unsolved regarding the control of triatomine vectors [4].

The subfamily Triatominae contains the main species responsible for the parasite transmission, particularly from the genera *Triatoma* (Laporte, 1832), *Panstrongylus* (Berg, 1879), and *Rhodnius* (Stål, 1859) [5]. *Rhodnius prolixus* is regarded as the main vector of *T. cruzi* in northern South America and Central America [6], and as a model organism in the study of insect physiology since Sir Vicent Wigglesworth published his seminal studies on the hormonal regulation of insect development [7]. In more recent years, the publication of *R. prolixus* genome and the establishment of molecular tools for functional studies (e.g. transcriptomes, *in situ* hybridization and RNA interference-RNAi) re-established this species as a great model system to study insect physiology, as well as vector-parasite interactions [8–11].

In the past ten years, comparative evolutionary and functional analysis provided evidence that genes containing small open reading frames (smORFs or sORFs, encoding < 100 aa polypeptides) constitute an unexplored and important part of eukaryotic genomes [12–14]. In this context, insects can represent excellent model organisms to study the role of these smORFs. For instance, the genome of *Drosophila melanogaster* encodes over 400 genes containing actively transcribed smORFs [15]. These smORF-containing genes are often transcribed from large mRNA precursors and can be polycistronic, e.g., more than one polypeptide can be encoded by a single mRNA.

The first polycistronic eukaryotic gene containing smORFs *mille-pattes* (*mlpt*) was identified in the genome of beetle *Tribolium castaneum* [16]. *mlpt* and its orthologs *polished rice* (*pri*) or *tarsal-less* (*tal*) play major roles during the embryogenesis of beetles, fruit flies, wasps and bugs [17–21]. Important roles of *mlpt/pri/tal* in the differentiation of epithelia, promotion of the development of trichomes, tarsus and gut precursors were also described[17,22]. Mechanistically, the conserved peptides of 10-13 amino acids encoded by the smORFs in the *mlpt* gene are involved in the control of N-terminal processing of the Shavenbaby (Sbv) transcription factor [22–26]. Importantly, mRNA levels of *mlpt/pri/tal* were shown to be modulated by ecdysone in *Drosophila melanogaster* [24,27]. However, strong evidence for a role of this gene in larval or nymphal development and moulting in other insects is still lacking.

Recently, multiple roles during embryogenesis were described for the single *mlpt* ortholog from *R. prolixus* (Rp-*mlpt*), and the existence of a putative new hemiptera-specific *mlpt* encoded peptide was also reported [20]. Thus, while the importance of Rp-*mlpt* during *R. prolixus* embryonic development is relatively well established, other functions of this gene in the kissing bug remain unknown.

In the present manuscript the postembryonic role of the smORF-containing gene Rp-*mlpt* was investigated in the kissing bug *R. prolixus*. As a hematophagous insect, *R. prolixus* digestion of haemoglobin generates haem, an important prosthetic group required for the activity of many enzymes, but also a highly reactive and toxic molecule. Elevated concentrations of haem are circumvented by several physiological mechanisms and most free haem in the midgut is aggregated into inert crystals of haemozoin, which are eliminated with the excreta and contribute to the dark-brown aspect of triatomine faeces [28,29].

Our results show that nymphs injected with Rp-mlpt double-stranded RNA (Rp-dsmlpt) do not moult, display delayed blood trafficking due to abnormal gut morphology and diminished peristalsis. These morphophysiological changes lead to a late accumulation of haemozoin in posterior midgut and a sharp reduction of total haem in the hindgut. Importantly, analysis of the influence of *Rp-mlpt* knockdown on the life cycle of *T. cruzi* revealed significant interference with parasite differentiation. Overall, this is the first report regarding the importance of a smORF-containing gene for post-embryonic development in a non-drosophillid insect, particularly in the moulting process and vector digestive physiology.

## Methods

### *Rhodnius prolixus* laboratory stock

*R. prolixus* bugs were kept in an incubator at 28 °C and 70 – 80% relative humidity. Different life cycle stages (eggs, nymphs and adult insects) were kept separately in plastic jars. Insects were fed through controlled exposure to rabbits and blood ingestion occurred through bites in the ears, as previously described [30]. Experimental protocols for this study were approved by the Ethics Committee on the Use of Animals from the Federal University of Rio de Janeiro (CEUA/UFRJ-Macaé), under licence code MAC032.

### *Trypanosoma cruzi* cell culture

Epimastigote forms of *Trypanosoma cruzi* Dm28c [31] were maintained at 28 °C in complete LIT medium [32] supplemented with 0.0025% hemin (Sigma-Aldrich), 10% inactivated fetal calf serum (Cultilab) and 10,000 U/L and 10 mg/L of penicillin and streptomycin (Sigma-Aldrich), respectively. The number of viable cells was determined by hemocytometer counts of motile cells. Epimastigotes grown to mid logarithmic phase with more than 90% viable cells were harvested for infection assays.

### Feeding and infection of experimental groups

Nymphs challenged with Rp-ds*mlpt* and ds*GFP* were fed by two distinct methods: (i) females were fed directly into the rabbit’s ear; (ii) females were fed in an artificial feeder at 37 °C with rabbit blood collected with a heparinized syringe.

For infection assays, rabbit blood was centrifuged during 10 min at 2000 g and 24 °C to separate plasma from erythrocytes. Then, the plasma was heat inactivated at 56 °C for 3 hours, while erythrocytes were resuspended and washed three times in phosphate-buffered saline, before being reunited with inactivated plasma. Simultaneously, *T. cruzi* epimastigotes were centrifuged during 10 min at 400 g and 16 °C and washed twice in phosphate-buffered saline. Parasites were then added to the recomposed blood to the concentration of 1 × 10^7^ cells/mL and this mixture was applied to the artificial feeder.

### RNA extraction, cDNA synthesis and RT-qPCR

Total RNA from fat body and midgut of *R. prolixus* was extracted at 1, 6 and 16 days after blood feeding (DAF). Alternatively, for *T. cruzi* infected triatomines, total RNA was extracted at 7, 16 and 21 DAF. RNA extraction was done using the TRIzol™ reagent (Invitrogen, Cat. No. 15596026), following the manufacturer-recommended procedures, concentration was estimated by spectrophotometry at 260 nm and purity was determined by 260/280 and 260/230 ratios, using a *Nanodrop 2000* spectrophotometer (Thermo Fisher Scientific). cDNA synthesis was carried with High-Capacity cDNA Reverse Transcription kit (Applied Biosystems, Cat. No. 4368814), following manufacturer’s instructions. Primers for Rp-*mlpt* and Rp-*eEF-1* have been previously described [20,33], while the primers for the ecdysone receptor (Rp-*EcR*) were designed for the present work, as follows: Rp-*EcR*-Fwd: TGTTGCGAATGGCTAGGAGG and Rp-*EcR*-Rev: CACCCATACCGGCCATACTG.

RT-qPCR amplifications were performed using *SYBR^®^ Green PCR Master Mix* (Applied Biosystems) on a *StepOnePlus*™ thermocycler (Applied Biosystems). All assays were performed with at least three technical replicates and a control without cDNA. Threshold cycles (Cq) were determined by default thermocycler parameters. Amplification efficiencies for each primer pair were determined using five-fold serial dilutions of fat body cDNA, as described [34]. Relative gene expression was accessed with the comparative 2^-ΔΔCq^ method [35], using Rp-*eEF-1* as reference of invariant expression, as previously reported [33,36].

### Gene knockdown via RNA interference

RNAi was performed as previously described [36]. Double stranded RNAs (dsRNAs) were generated with T7 Megascript Kit (Ambion, Cat. No. AM1334). Primers for Rp-*mlpt* dsRNA were T7-Rp-*mlpt*-Fwd: ggccgcggAATACGCTCGATCCTACCGG and T7-Rp-*mlpt*-Rev: cccggggcTGGTCTGTATTAATCCC. Nucleotides in lowercase letters in the primers were used for a second PCR reaction to add T7 promoter sites at the extremities. The green fluorescent protein (GFP) gene sequence was used as a template for the synthesis of an unrelated control dsRNA, using the following primers: T7-GFP-Fwd: taatacgactcactatagggACGTAAACGGCCACAAGTTCAGCGTGTC and T7-GFP-Rev: taatacgactcactatagggTCACGAACTCCAGCAGGACCATGTGATC (lower case letters corresponds to the T7 adaptor sequence). 4 μg of dsRNA were injected in unfed nymphs for each target gene and the insects were fed with blood 24 hours after the injection. Silencing efficiency was assessed through RT-qPCR, using the comparative 2^-ΔΔCq^ method, as described above. Alternatively, the same amount of dsRNA was injected in *R. prolixus* nymphs at 3, 7 and 14 DAF.

### Immunofluorescence microscopy

For immunofluorescence staining, insects were dissected, and tissues were collected and fixed at room temperature with 4% paraformaldehyde for one day, followed by staining with phalloidin conjugated to Alexa Fluor 488 (1:100 dilution; Sigma-Aldrich) and DAPI (1:500 dilution; Sigma Aldrich) for 30 min in phosphate-buffered saline with Tween-20 1% (v/v), as previously described [37]. Images were acquired using a Leica M205 stereoscopic microscope.

### Haemozoin extraction

Haemozoin was extracted from the midgut of *R. prolixus* as previously described [38]. Final sediments were solubilized in 0.1 M NaOH and haem amount was determined by spectrophotometry at 400 nm in a *UVmini-1240* spectrophotometer (Shimadsu).

### *T. cruzi* quantification in *R. prolixus* hindgut

Infected insects were dissected, and hindgut samples were homogenised in 100 μL of phosphate-buffered saline. The number of parasites was determined by counting in a Neubauer haemocytometer. *T. cruzi* epimastigotes were differentiated from metacyclic trypomastigotes based on their characteristic morphologies and motility patterns. Intermediate stages were considered as epimastigotes.

### Statistical Analysis

For normally distributed data, comparisons between means of two independent groups were performed with the unpaired t-test, and multiple comparisons were done with one-way analysis of variance (ANOVA), followed by Tukey post-test. For nonparametric data, pairwise comparisons were performed with the Mann-Whitney test, while multiple comparisons were done with the Kruskal-Wallis test, followed by Dunn’s post-test. Results are expressed as group means ± standard deviations. Differences were considered significant whenever p ≤ 0.05. Graphpad Prism 6 software (v. 6.01) was used for statistical analysis.

## Results

### Rp-*mlpt* expression is differentially regulated by blood feeding and its knockdown alters blood digestion in fifth instar nymphs of *Rhodnius prolixus*

To understand the role of Rp-*mlpt* during the development of *R. prolixus*, we have first examined the relative expression of its mRNA on all nymphal stages, and no significant increase or decrease in mRNA levels was observed along the development (supplementary figure S1). Notably, Rp-*mlpt* is differentially regulated after blood feeding in several organs in adulthood, being upregulated in the fat body and downregulated in the midgut and ovaries (figure 1).

**Figure 1.**
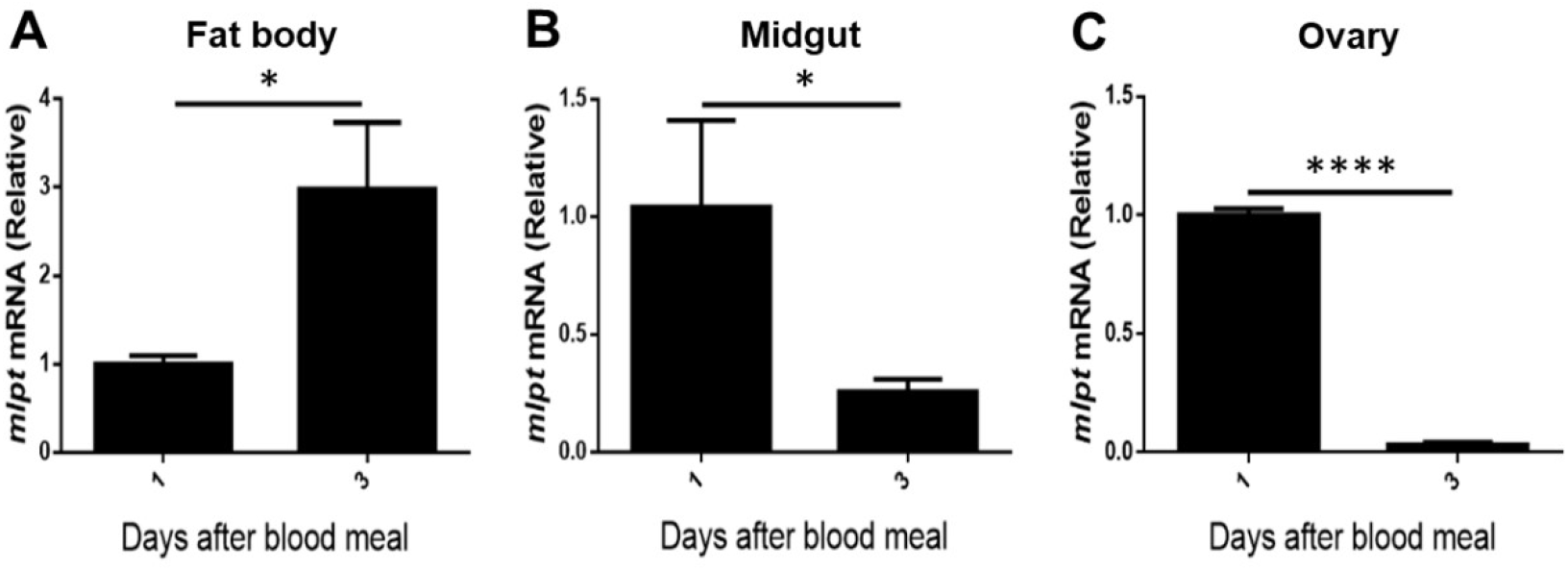
Rp-*mlpt* is differentially regulated after blood feeding in several organs in adult *R. prolixus*. Rp-*mlpt* relative mRNA expression determined by RT-qPCR in fat body (A), midgut (B) and ovary (C) of blood-fed adult females at 1 and 3 days after feeding. Rp-*eEF-1* mRNA was used as a reference of invariant expression (endogenous reference gene). Data are represented as the mean ± standard deviations of three independent experiments and expressed as fold change of calibrator expression. Statistical analysis was performed using unpaired student-t tests. * p ≤ 0.05; **** p ≤ 0.0001.

**Supplementary figure 1.**
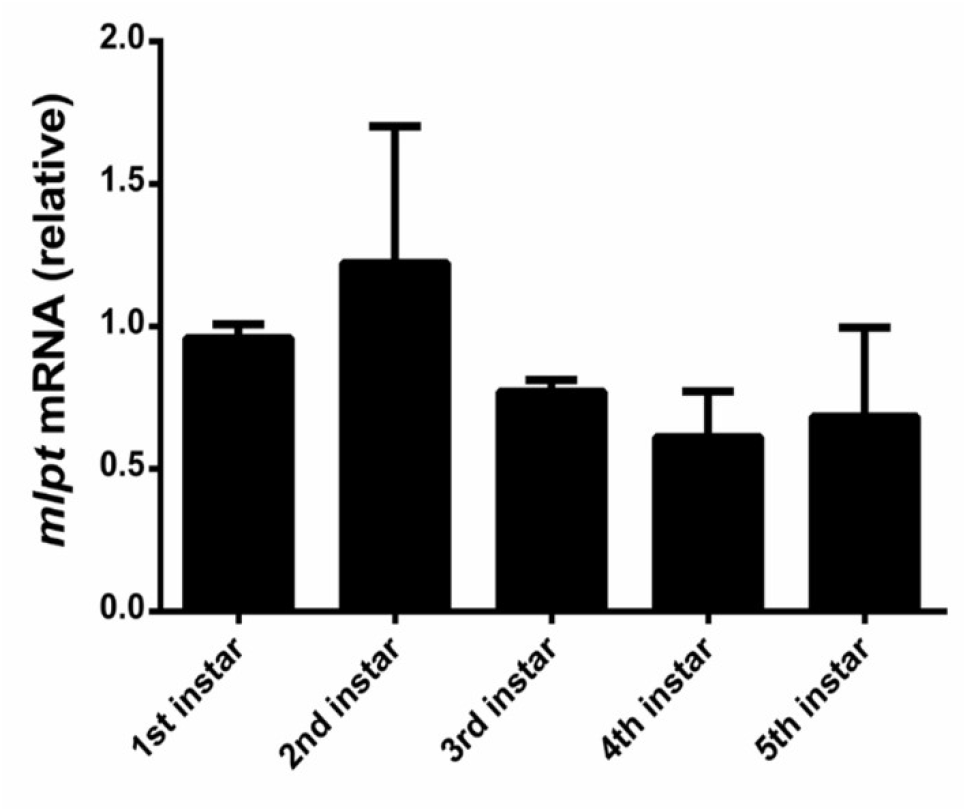
Rp-*mlpt* is expressed during all nymphal stages of *R. prolixus*. Rp-*mlpt* relative mRNA expression along nymphal development determined by RT-qPCR. Rp-*eEF-1* mRNA was used as a reference of invariant expression (endogenous reference gene). Data are represented as the mean ± standard deviations of three independent experiments and expressed as fold change of calibrator expression. Statistical analysis was performed using ANOVA.

To further investigate the role of Rp-*mlpt*, unfed fifth instar nymphs were injected with double-stranded RNA (dsRNA) to knockdown its expression. The challenge with Rp-ds*mlpt* (double-stranded Rp-*mlpt*) was more responsive in the fat body, while there was no significant difference in mRNA expression in the posterior midgut (supplementary figure S2*a*). In the fat body, the expression of the Rp*-mlpt* gene in the Rp-ds*mlpt* group decreased within one day after blood feeding (DAF), and remained at low levels until at least six DAF (supplementary figure S2b).

**Supplementary Figure 2.**
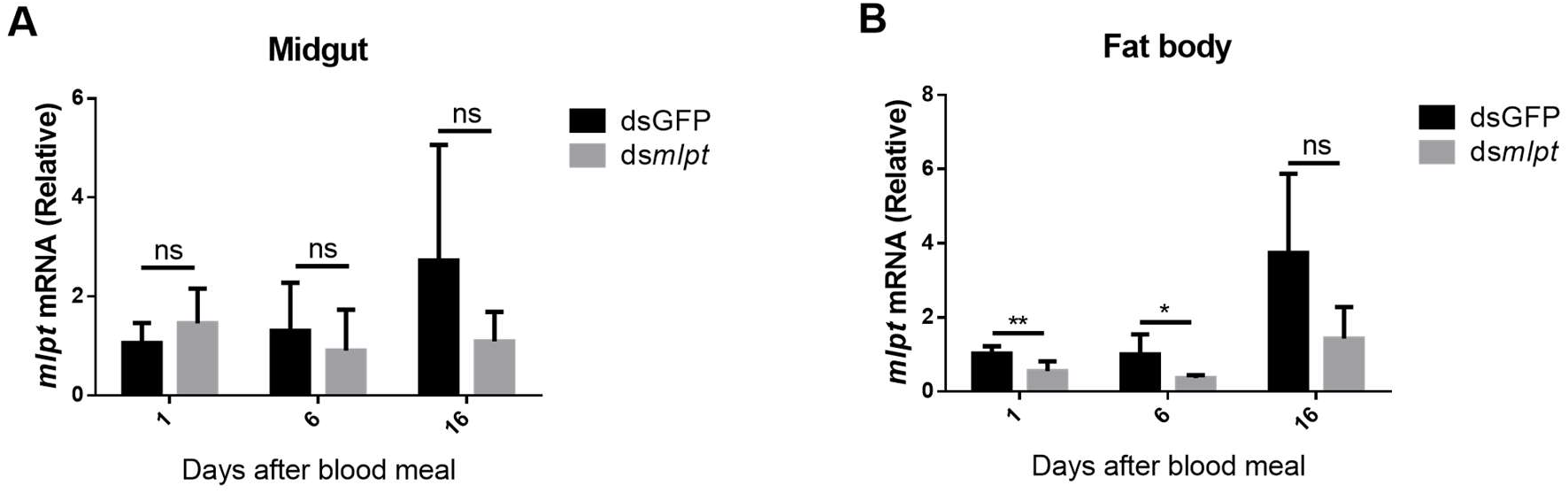
Relative expression of Rp-*mlpt* mRNA in *R. prolixus* after injection with Rp-ds*mlpt*. A: Rp-*mlpt* relative mRNA expression in the midgut of 5th instar blood fed at 1, 6 and 16 DAF. B: Rp-*mlpt* relative mRNA expression in the fat body of blood fed 5th instar at 1, 6 and 16 DAF. Rp-*eEF-1* was used as a reference of invariant expression and ds*GFP* RNA was alternatively injected in order to provide a negative control of RNA interference. Data are represented as means ± standard deviations of three independent experiments and expressed as fold change of calibrator expression. Statistical analysis was performed using unpaired student-t tests. ns: non-significant (p > 0.05); * p ≤ 0.05; ** p ≤ 0.01.

Next, we examined the phenotypic alterations that resulted from the Rp-*mlpt* RNAi knockdown. Midgut external morphology showed no discernible difference between control and Rp-ds*mlpt* groups up to 6 DAF (figure 2*a*,*b*). In contrast, after 16 DAF, hindgut morphology and function were severely affected in the Rp-ds*mlpt* group. The hindgut was abnormally increased in the group treated with Rp-ds*mlpt*, when compared to controls (figure 2*a*,*b*). In addition, a decrease in anterior midgut peristaltic movements in Rp-ds*mlpt* was evident at 16 DAF (supplementary video 1,2), suggesting that traffic of ingested blood along the digestive tract was reduced after RNAi. The inspection of tissue organisation by fluorescence microscopy revealed that the microfilament architecture was unaffected at 6 DAF, but microfilaments were considerably disrupted at 18 DAF in the posterior midgut (PM) of Rp-ds*mlpt* (figure 2b). This observation revealed that the muscle layers surrounding the midgut epithelium were seriously compromised due to Rp-*mlpt* knockdown. This disruption in visceral muscular layers was not generalised, as muscle layers from the salivary gland were not visibly altered after 18 days DAF (supplementary figure S3).

**Figure 2.**
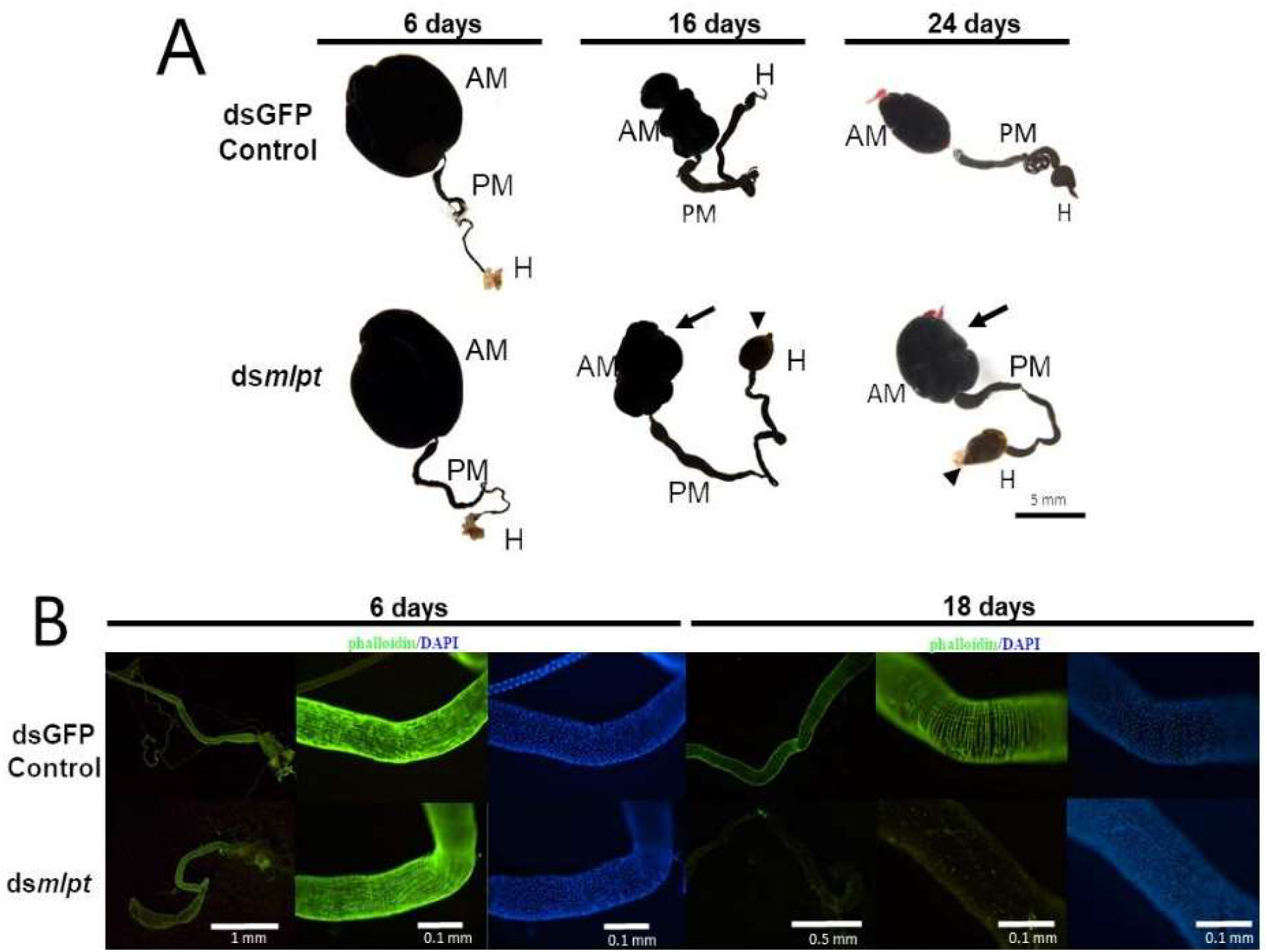
RNAi silencing of Rp-*mlpt* gene affects posterior midgut and hindgut morphology and function. A: Overall digestive tract morphology of ds*GPF* (control) and Rp-ds*mlpt* RNAi (*dsmlpt*) 6, 16 and 24 DAF. AM: anterior midgut; PM: posterior midgut; H: hindgut. Arrows indicate increase in size of the anterior midgut and arrowheads indicate increase in size of hindgut for Rp-*mlpt* silenced insects. B: Nuclear (DAPI - blue) and filamentous actin (phalloidin-Alexa 488 - green) fluorescent staining of Rp-ds*mlpt* and ds*GFP* posterior midgut. Phalloidin and DAPI staining were similar at 6 DAF among Rp-ds*mlpt* and ds*GFP* midguts, while irregular phalloidin staining was evident at 18 DAF for Rp-ds*mlpt*.

**Supplementary figure 3.**
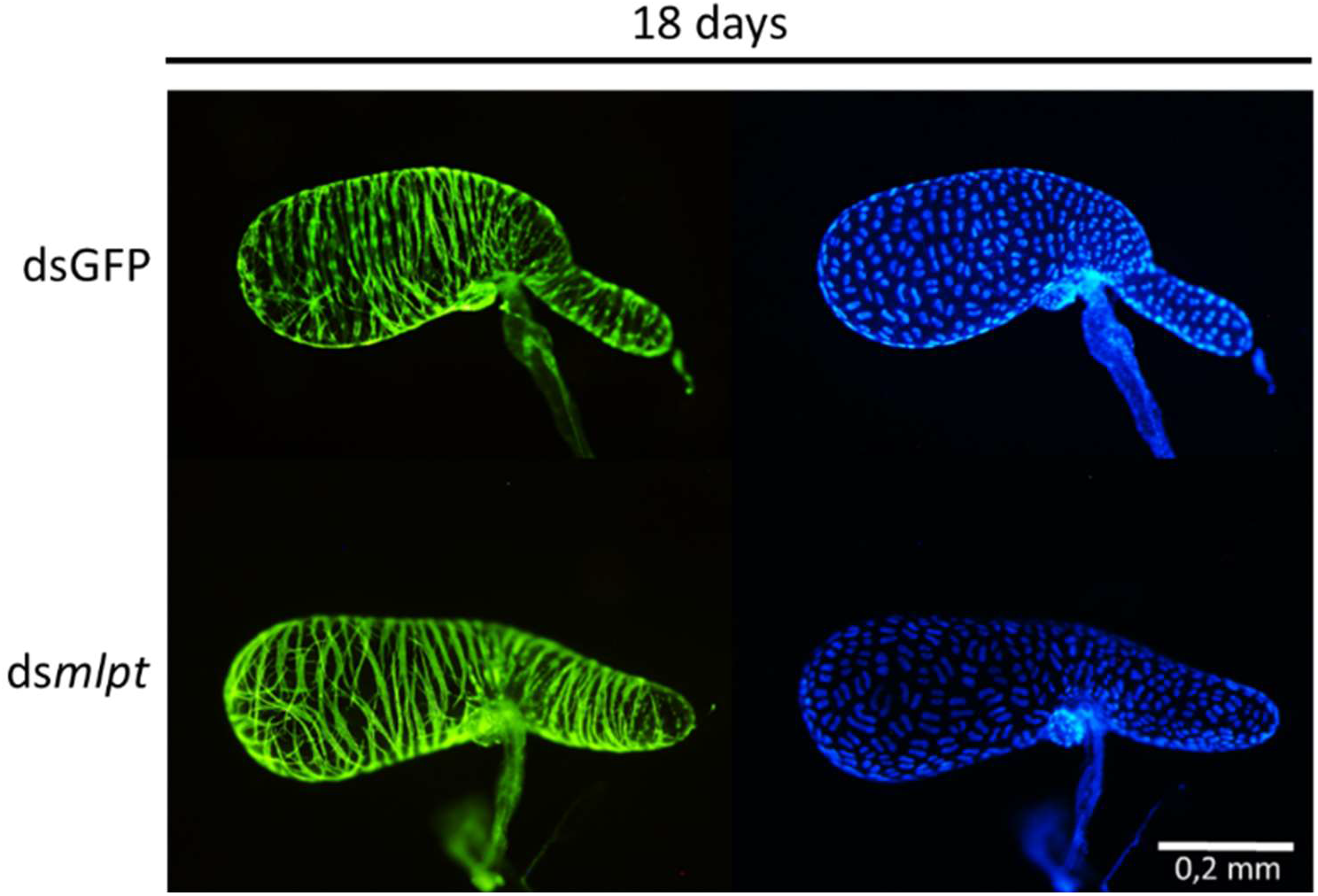
RNAi silencing of the Rp-*mlpt* gene does not interfere with nuclei and muscle layer organization of *R. prolixus* salivary glands. Nuclear (DAPI - blue) and filamentous actin (phalloidin-Alexa 488 - green) fluorescent staining of Rp-ds_mlpt_ and ds*GFP* dissected salivary glands. Phalloidin and DAPI staining were similar at 18 DAF for both Rp-ds*mlpt* and ds*GFP* controls.

To determine if blood intake is affected by Rp-*mlpt* knockdown, nymphs had their body weight determined shortly after blood feeding. In addition, the body weight of the nymphs was monitored for 16 days after the blood meal. Insect weight was similar in control and nymphs injected with Rp-ds*mlpt*, indicating that blood intake and excretion over time are not affected by Rp-*mlpt* knockdown (figure 3a).

**Figure 3.**
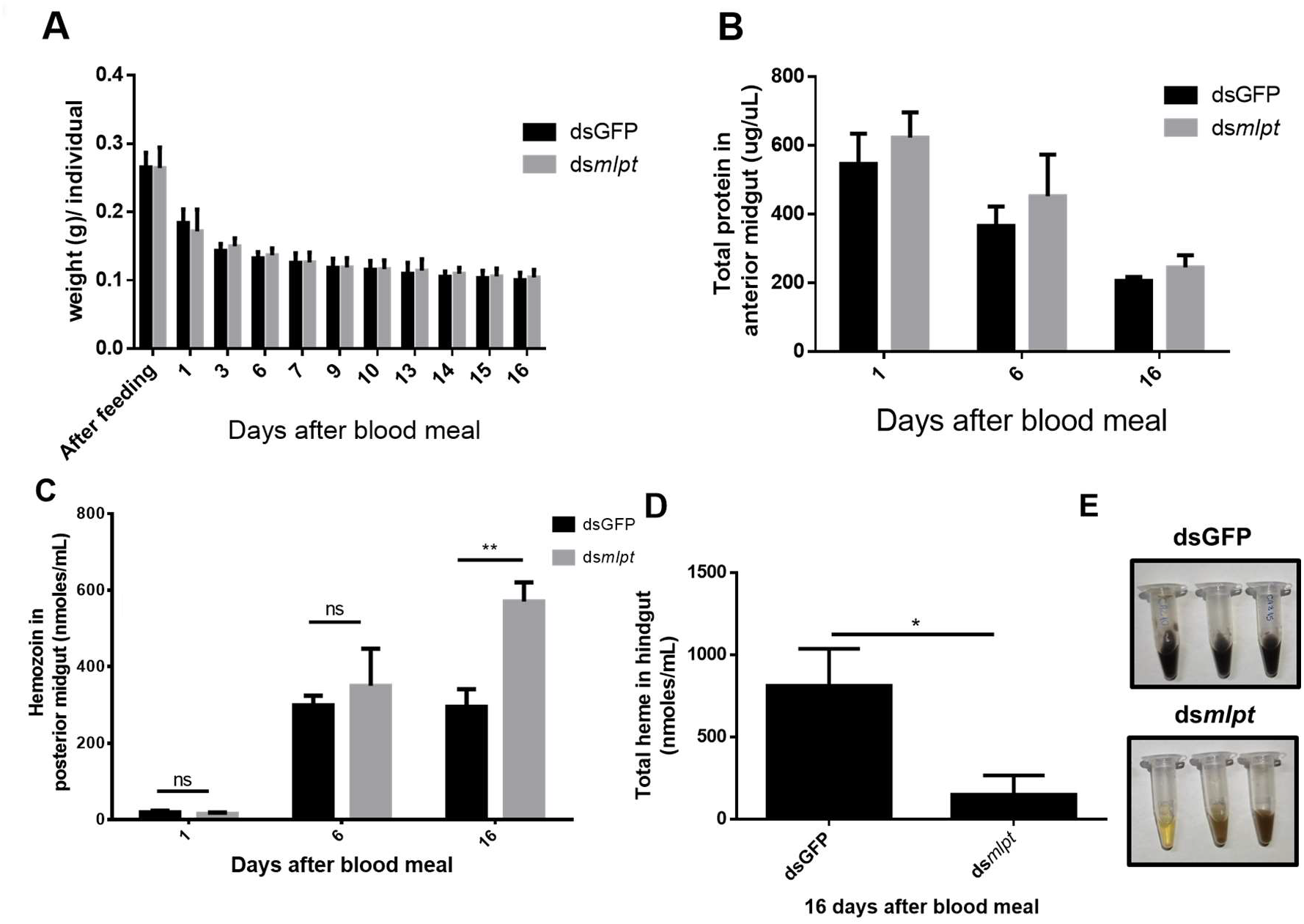
RNAi silencing of Rp-*mlpt* gene does not affect blood digestion, but modifies the contents of the posterior midgut and hindgut. A, B: Rp-*mlpt* silencing does not affect body weight (A) and total protein content in the anterior midgut (B) until 16 DAF. Protein content in (B) was assessed with the BCA method. C: A significant increase in haemozoin content is observed in the posterior midgut (PM) of Rp-ds*mlpt* challenged insects at 16 DAF. D: Total haem (unpolymerized haem + haemozoin) is significantly reduced in the hindgut at 16 DAF of Rp-ds*mlpt*. E: The visual aspect of hindgut contents at 16 DAF is distinct between Rp-ds*mlpt* challenged insects and the ds*GFP* control group. Data in graphs were analysed with unpaired t-tests. ns: non-significant (p > 0.05); * p ≤ 0.05; ** p ≤ 0.01.

Next, we investigated the digestion route of blood haemoglobin and products derived from its breakdown. Total protein levels measured in the anterior midgut (AM) in the Rp-ds*mlpt* group and in the control were equivalent at 1, 6 and 16 DAF (figure 3b), indicating that haemoglobin digestion was occurring normally in both groups. Interestingly, until the 6th DAF, both experimental groups produced haemozoin crystals in the same proportion (figure 3c), but on the 16th DAF the Rp-ds*mlpt* group had accumulated more haemozoin in the PM (figure 3c). When total haem in the hindgut was quantified, a concentration five times lower was determined for the Rp-ds*mlpt* group when compared to the control. Additionally, we noted that the hindgut content from insects challenged with Rp-ds*mlpt* presented a yellowish-brown colour, in contrast to the usual dark-brown colour observed in the control group, probably reflecting the reduction in haem content observed in this compartment (figure 3d). These results indicate that the haemozoin produced in the PM does not reach the hindgut, probably due to the effect of Rp-ds*mlpt* on midgut muscle layers and, consequently, on its peristalsis. Overall, these data demonstrate severe digestive defects in Rp-ds*mlpt* nymphs, which are mainly restricted to the latest stages of digestion (16 DAF). Thus, the *R. prolixus* Rp-*mlpt* gene is essential for proper blood traffic and digestion during nymphal development.

### Rp-*mlpt* knockdown inhibits moulting

To further investigate the roles of Rp-*mlpt* in the life cycle of *R. prolixus*, Rp-*dsmlpt* RNA was injected into unfed fifth instar nymphs. Notably, all nymphs injected with Rp-ds*mlpt* RNA were unable to moult to adulthood (figure 4), indicating that Rp-*mlpt* is critically important for moulting in *R. prolixus*. To investigate the time window of Rp-*mlpt* requirement for moulting, Rp-ds*mlpt* was also injected into fifth instar nymphs at different DAF (3rd, 7th, 14th DAF) and the impacts on moulting to the next stages were analysed. All fifth instar nymphs injected with Rp-ds*mlpt* at the 3rd and 7th DAF were also unable to moult, while all nymphs injected at the 14th DAF were fully able to moult to the adult stage (figure 4). These results show that the requirement of Rp-*mlpt* for nymphs to moult is restricted to the period between 7th and 14th DAF.

**Figure 4.**
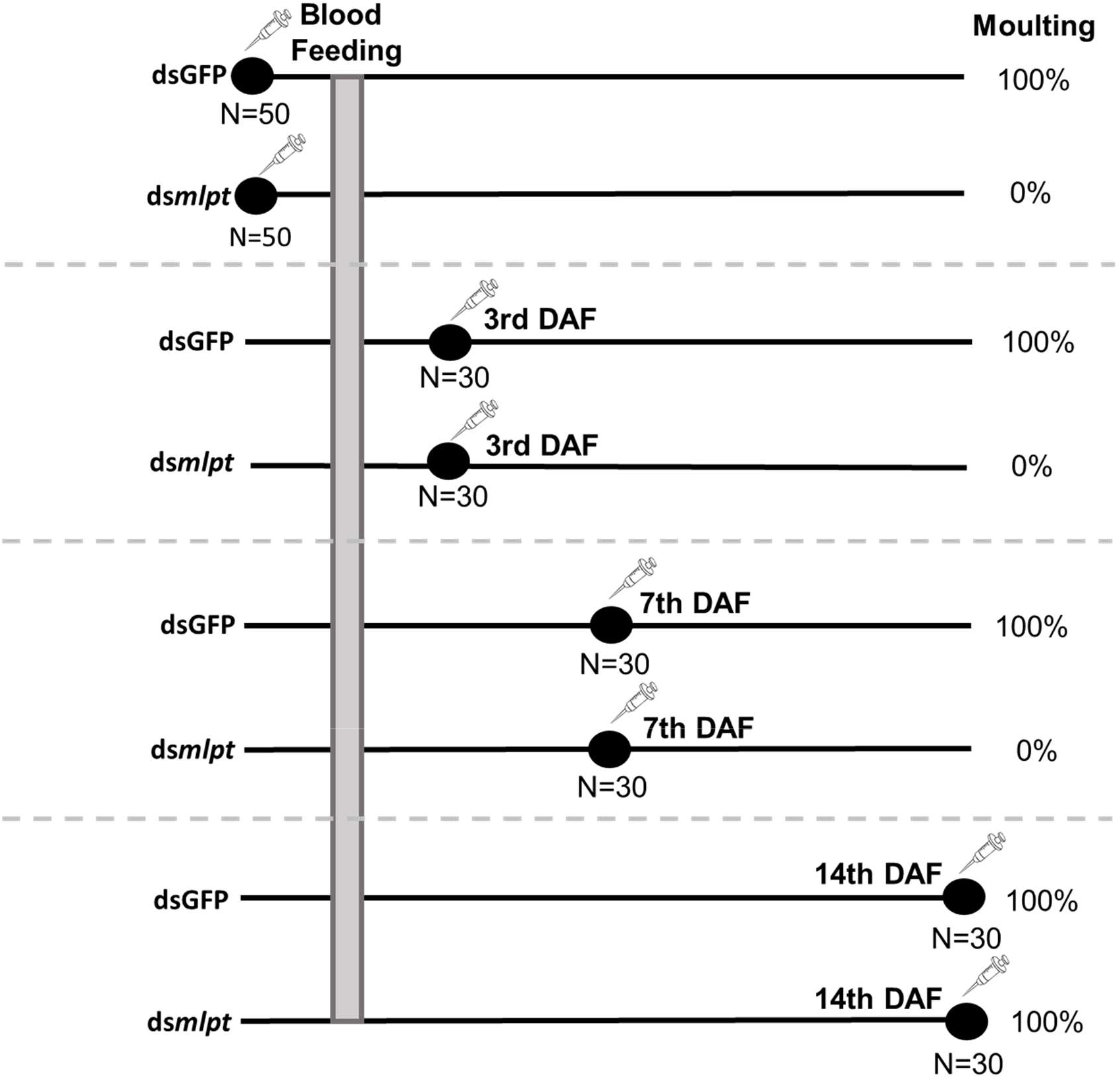
Rp-*mlpt* RNAi knockdown blocks moulting. Moulting after dsGPF or Rp-ds*mlpt* injection into fifth instar nymphs. Moulting is completely blocked in all nymphs injected with Rp-ds*mlpt*, except for the 14th DAF, which moult normally (n=30). In the control group, all individuals moult between 16 and 28 DAF, regardless of the time of ds*GFP* injection.

To reveal whether moulting dependence on Rp-*mlpt* expression occurs only in the transition from the fifth nymphal to the adult stage, unfed fourth instar nymphs were also injected with ds*GFP* and Rp-ds*mlpt*. Once more, all Rp-*mlpt* silenced insects failed to moult to the fifth instar (n=7), while control insects moulted normally (n=10), suggesting that every moult during *R. prolixus* development relies upon Rp-*mlpt* expression. The impact of Rp-*mlpt* knockdown on moulting cannot be attributed to the downregulation of the ecdysone receptor (EcR) expression, as the relative expression of Rp-*EcR* mRNA in the midgut and fat body showed no significant difference between the two experimental groups (supplementary figure S4a,b).

**Supplementary figure 4.**
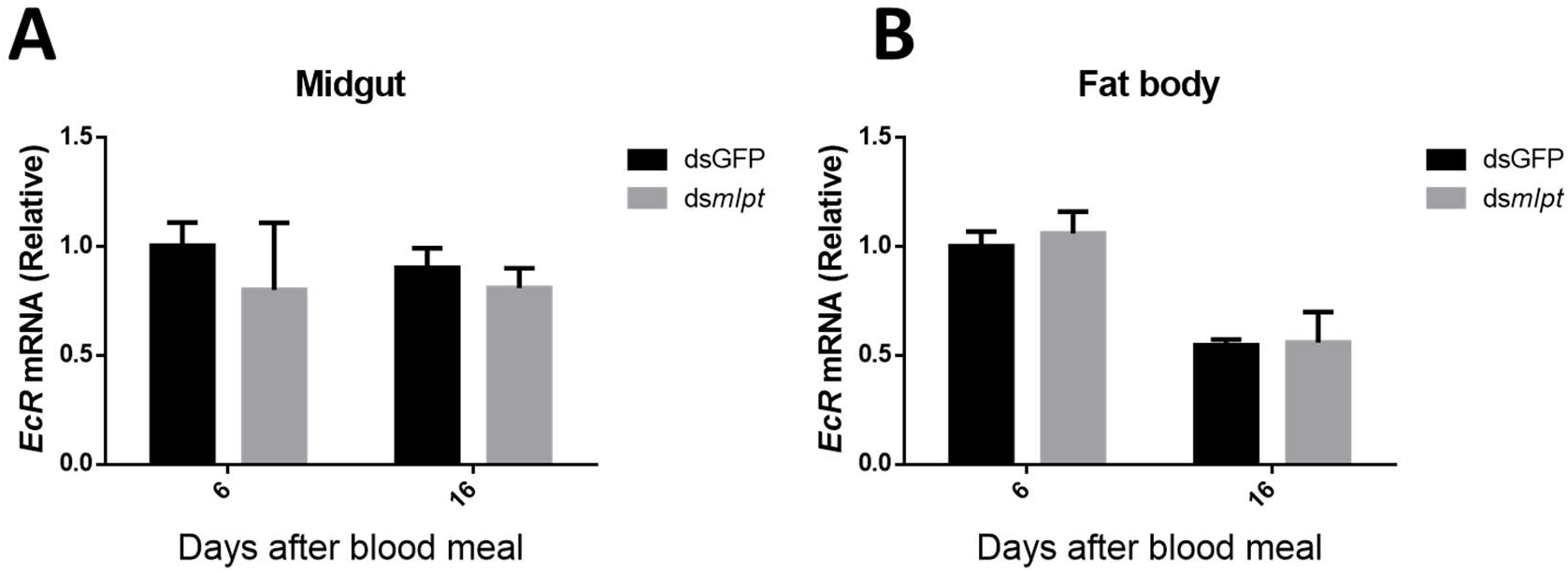
Rp-*mlpt* silencing does not affect ecdysone receptor (*EcR*) transcript levels. A, B: Relative expression of Rp-*EcR* mRNA in the midgut (A) and fat body (B) of control and Rp-ds*mlpt* injected females at 6 and 16 DAF. Rp-*eEF-1* was used as reference of invariable expression and ds*GFP* RNA was alternatively injected to provide a negative control of RNA interference. Data are represented as the mean ± standard deviations of three independent experiments and expressed as fold change of calibrator expression. Statistical analysis was performed using unpaired student t-tests.

### Rp-*mlpt* is essential for *T. cruzi* metacyclogenesis during nymphal development

The hormone ecdysone is essential for the differentiation from proliferative epimastigotes to the metacyclic trypomastigote infective stage, a process known as metacyclogenesis, which occurs in the hindgut and is dependent on the microenvironmental conditions of the intestinal lumen [39]. Since Rp-*mlpt*, a putative target of ecdysone, is essential for moulting during nymphal stages, we hypothesised that it could interfere with *T. cruzi* proliferation and metacyclogenesis along the insect digestive tract. Upon *T. cruzi* artificial infection, both experimental groups (ds*GFP* and Rp-ds*mlpt*) presented the same weight pattern and similar total protein amount in the AM over time (figure *5a,b*), as noted for non-infected nymphs (figure *3a,b*). As previously observed for non-infected animals (figure 3c,d), the Rp-ds*mlpt* group infected with *T. cruzi* also accumulated more haemozoin at later time points then the ds*GFP* controls (figure 5*c*).

**Figure 5.**
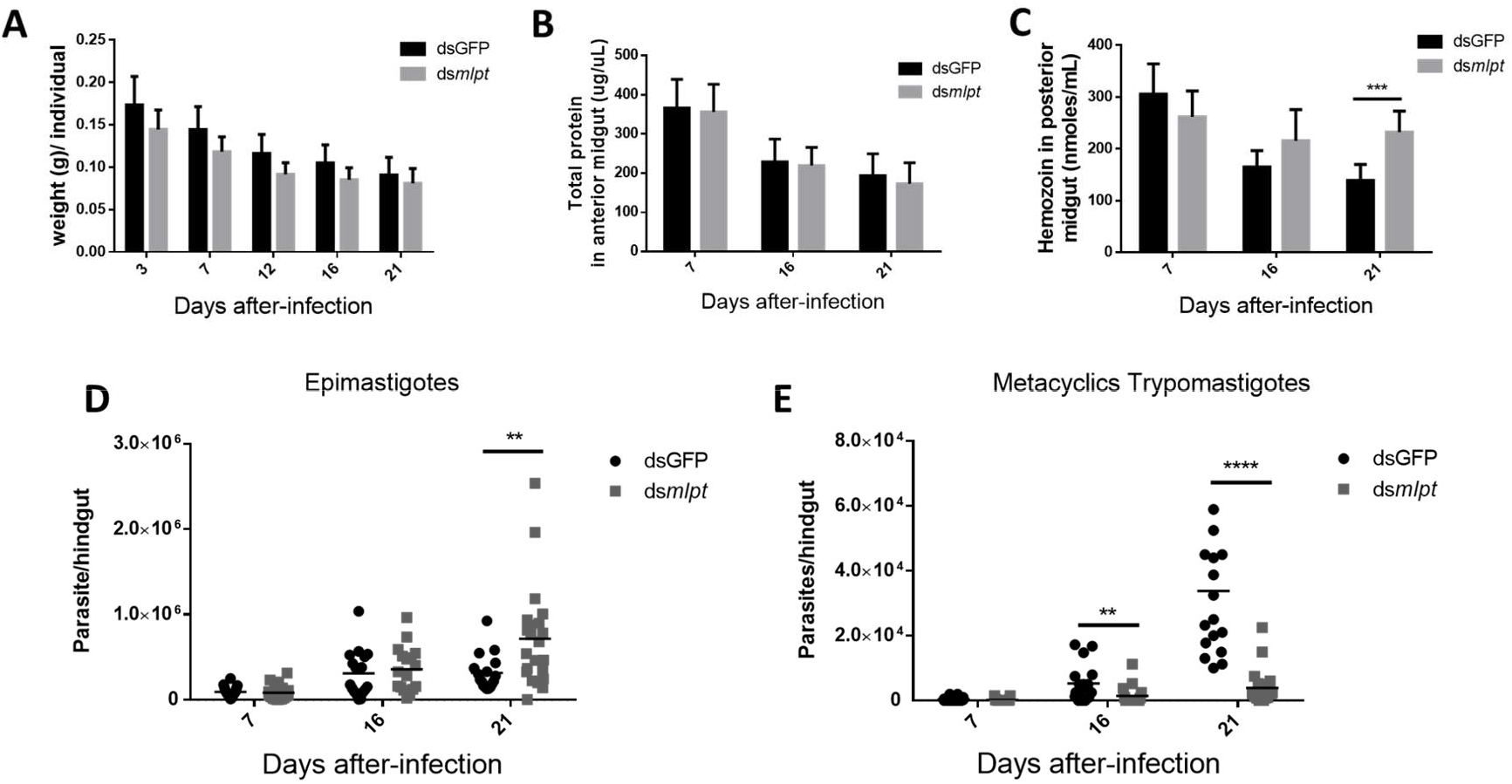
Rp-*mlpt* is essential for *T. cruzi* metacyclogenesis during nymphal stages. A, B: Rp-*mlpt* silencing does not affect body weight (A) and total protein content in the anterior midgut (B) of *T. cruzi* infected nymphs. Protein content in (B) was assessed with the BCA method. C: A significant increase in haemozoin content in PM is observed in Rp-ds*mlpt* challenged insects 21 days after feeding with infected blood. D, E: Influence of Rp-*mlpt* silencing on *T. cruzi* life cycle as revealed by total number of parasites. Epimastigotes (D) and metacyclic trypomastigotes (E) in the hindgut of control ds*GFP* and Rp-ds*mlpt* insects at 7, 16 and 21 DAF with infected blood. A significant increase of total parasites was observed for Rp-ds*mlpt* at 21 DAF, while significant reduction in the number of metacyclic trypomastigotes was observed for Rp-ds*mlpt* at 16 and 21 DAF. Data in A, B and C graphs were analysed with unpaired t-tests; data in D and E graphs were analysed with Mann-Whitney tests. ** p ≤ 0.01; *** p ≤ 0.001; p ≤ 0.0001.

To investigate the effects of Rp-*mlpt* knockdown in *T. cruzi* life cycle, parasite load and differentiation status were analysed at 7, 16 and 21 DAF. While no significant difference was observed in the total number of parasites in the hindguts of Rp-ds*mlpt* nymphs at 7 and 16 DAF, at 21 DAF more epimastigotes were present in the group challenged with Rp-ds*mlpt* (figure 5d). In the case of the infective, metacyclic trypomastigotes, after 16 DAF (before moulting) and 21 DAF (after moulting in the ds*GFP* group) the number of these forms in the hindgut of Rp-ds*mlpt* nymphs was drastically reduced, when compared to control ds*GFP* nymphs and adult insects (figure 5e). These data show that the interference with Rp-*mlpt* expression in the triatomine vector has a profound impact on *T. cruzi* differentiation into infective forms.”

We further observed the population of parasites collected from the hindguts from both groups. In the ds*GFP* control group, we observed the typical morphologies of epimastigotes (arrowheads in figure 6*a*,*b*) and metacyclic trypomastigotes (arrows in figure 6*a*,*b*). In the Rp-ds*mlpt* group, while epimastigote forms were more abundant and trypomastigotes scarce, forms resembling intermediate differentiation stages were found in relative abundance, as highlighted in figure 6*c*,*d*, suggesting that the parasites started the differentiation process, but failed to accomplish it.

**Figure 6.**
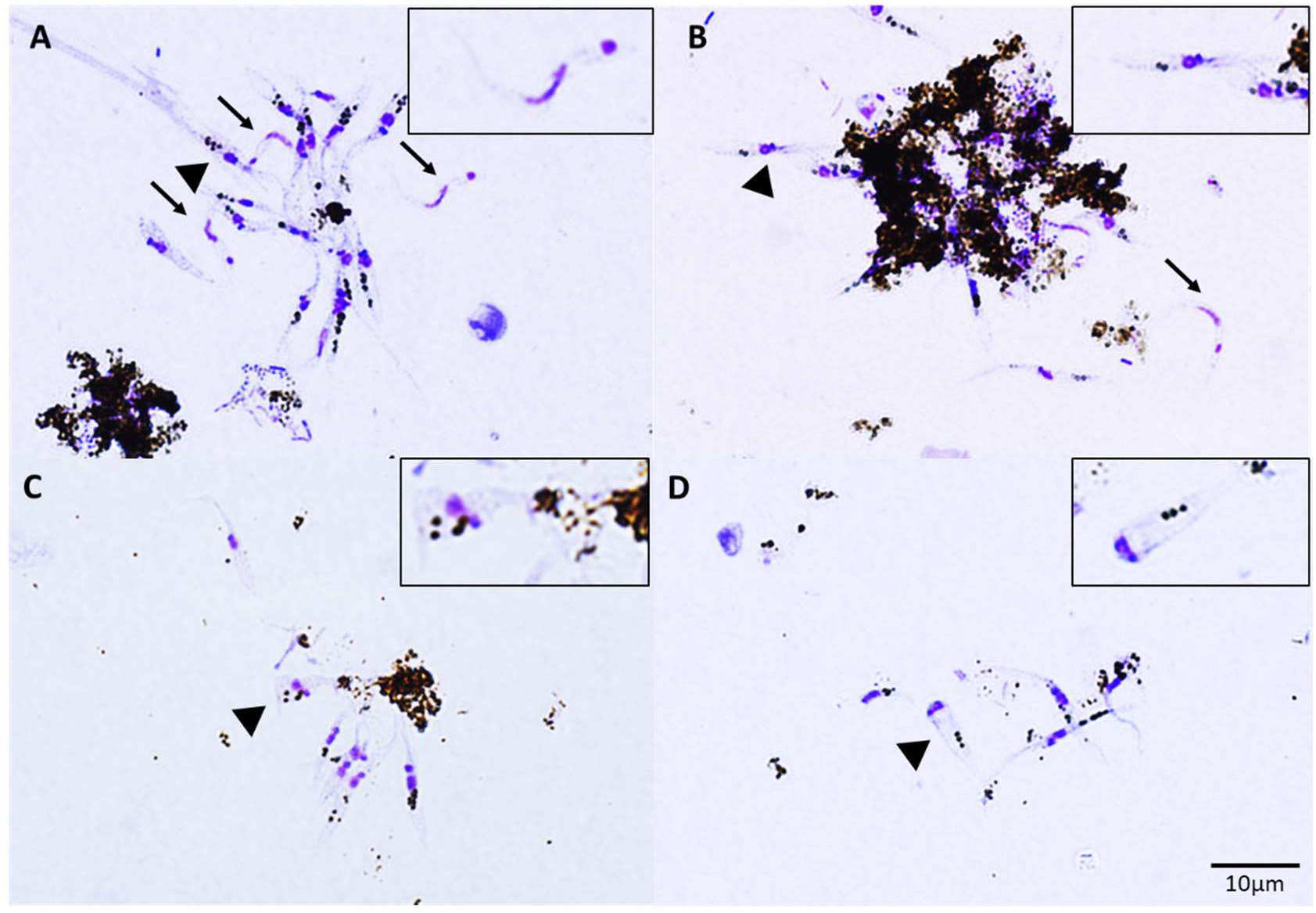
Rp-*mlpt* silencing interferes with *T. cruzi* metacyclogenesis. Microscopic analysis of hindgut contents from control (dsGFP) and Rp-ds*mlpt* challenged nymphs of *R. prolixus* 14 days after feeding with *T. cruzi* infected blood. Hindgut contents were stained with fast-panoptic and observed in brightfield, under 100× oil immersion lens. Nuclei and kinetoplast are stained in dark blue. A, B: representative imagens of control (dsGFP) parasites. C, D: representative images of Rp-ds*mlpt* challenged parasites. Arrows point to typical metacyclic trypomastigotes, while arrowheads indicate the selected specimens highlighted in insets: metacyclic trypomastigotes in A, epimastigote in B and intermediate forms in C and D. Darker brown granules in parasites probably correspond to endocytic reservosome compartments present in epimastigote and intermediate forms but absent from mature trypomastigotes.

## Discussion

It is currently known that hundreds of genes containing smORFs are present in insect genomes and may be evolutionary associated with the evolution of more complex genes and new cellular functions [40]. However, due to a plethora of technical difficulties, ranging from *in silico* prediction to the establishment of functional assays, the vast majority of these smORFs are still insufficiently investigated. Here we report that one gene encoding smORFs, Rp-*mlpt*, is essential for the integrity of longitudinal and circular intestinal muscle layers and, consequently, for efficient peristalsis. *R. prolixus* nymphs challenged with Rp-ds*mlpt* show dramatic changes in gut morphology and reduced traffic of blood digestion metabolites from posterior midgut to hindgut, which ultimately interferes with the natural cycle of the pathogen *T. cruzi*.

At least two technical limitations must be reported for the analysis and interpretation of Rp-*mlpt* RNAi results. First, the injection of a dsRNA in the haemolymph of insects leads to a systemic knock-down of gene expression [41,42], thus, it is not clear where Rp-*mlpt* encoded peptides are produced and act in *R. prolixus* wild-type nymphs. Second, the small size of smORF-encoded peptides precludes the development of antibodies and their applications in western-blots and immunostainings as well their detection via common proteomic techniques.

### Rp-*mlpt* is required for proper peristalsis, digestion and excretion during nymphal development

We have first examined the differential regulation of Rp-*mlpt* mRNA expression upon blood feeding. While Rp-*mlpt* expression is upregulated in the fat body three days after blood feeding, its expression is severely downregulated in ovaries and midgut. The upregulation of Rp-*mlpt* in the fat body is noteworthy, since this tissue is highly dynamic and secretive, and exerts pivotal metabolic influence over the whole organism [43]. Thus, the Rp-*mlpt* peptides produced and secreted by the fat body in response to blood feeding might have a broad impact in *R. prolixus* physiology. The results presented here suggest that Rp-*mlpt* encoded peptides are essential to synchronise digestion and excretion between blood meals.

In *R. prolixus*, the RNAi mediated silencing of Rp-*mlpt* does not affect blood digestion in the anterior midgut significantly until the 16th DAF (figure 3b). However, a notable decrease in midgut peristalsis was observed in later periods for the Rp-ds*mlpt* group, which probably occurred due to the disorganisation of the intestinal muscular layers, as revealed by phalloidin staining of actin filaments (figure 2b). In this context, the unusual accumulation of haemozoin crystals in the posterior midgut (figure 3c) contrasts with the amount of total haem found in the hindgut (figure 3d). It is conceivable that this phenomenon occurs due to the defective intestinal peristalsis, which leads to haemozoin accumulation in the midgut and diminished traffic to the hindgut. The light yellowish-brown coloured content observed in the hindgut also reflects the reduced content of haemozoin found in this segment (figure 3e). Both anterior midguts and hindguts of Rp-*mlpt* silenced individuals are visibly engorged (figure 2a), probably due to the reduced peristalsis leading to the accumulation of ingested blood in the anterior segment. It is possible that excretion through the hindgut begins to be impaired after the 6th DAF, culminating, at later times (e.g. 16th DAF), in a clogged and distended hindgut, whose luminal content may be composed of residues derived from the hindgut and haemolymph fluid.

It is well known that the constant turnover of the insect’s gut tissue and peristaltic movement are important for digestion and excretion [44,45]. In *D. melanogaster*, the hormone ecdysone has been shown to be essential for activating the production of *mlpt* peptides within enteroendocrine cells in the adult intestine, which, in turn, triggers the activation of the Shavenbaby (Svb) transcription factor to maintain intestinal stem cells [37]. It is also known that a set of *Drosophila* enteroendocrine cells are required for efficient peristalsis and secretion [46]. In this context, if Rp-*mlpt* is also essential for intestinal cell renewal in *R. prolixus*, as described for *Drosophila*, it is possible that that the intestinal damage observed in nymphs challenged with Rp-ds*mlpt* may be connected to a dysfunction in enterocytes, which could indirectly result in severely impaired muscle microfilament network, defective peristalsis and enlarged intestine segments.

### Rp-*mlpt* expression between 7 and 14 DAF is essential for moulting

Since feeding and moulting are known to be functionally linked events in insect physiology [47], it is conceivable that the aforementioned deleterious effects may promote interference with moulting and ecdysis. Indeed, here we show that Rp-*mlpt* is important for the proper moulting of *R. prolixus* nymphs.

After silencing the Rp-*mlpt* gene, there is a critical impairment in the moulting from the fifth instar nymph to the adult stage, as well as from the fourth to the fifth instar nymph. Interestingly, Rp-ds*mlpt* injections at 3rd and 7th DAF lead to the same phenotype, the absence of moulting, while injection of Rp-ds*mlpt* at 14th DAF does not impair moulting, indicating that this biological phenomenon requires the expression of Rp*-mlpt* expression between 7th and 14th DAF. In *R. prolixus*, moulting is coordinated and synchronised whenever an adequate blood meal is taken by a nymph [48]. This crucial physiological process is strictly regulated by the neuroendocrine system, which involves neuropeptides and their receptors in target cells [47]. About 41 putative neuropeptide precursor genes have already been identified in *R. prolixus*, but only little is known about their functions in the neuroendocrine system [9,49]. Interestingly, Wulff and colleagues [50,51] documented that the three different isoforms of orcokinins neuropeptides (RhoprOKA, RhoprOKB and RhoprOKC) are associated with the regulation of post-embryonic development and digestive physiology. Injection of *RhoprOKA* dsRNA inhibited moulting, while silencing of the *RhoprOKB/C* isoforms lead to delayed moulting [51]. Since Rp-*mlpt* silencing effects described here are similar to those of *RhoprOKA* silencing, both genes appear to be important integrators of nutritional status with moulting. Whether Rp-*mlpt* encoded peptides act downstream or upstream of *RhoprOKA*, or if both genes act in parallel signalling pathways to regulate moulting remains to be investigated.

### Phenotypic analysis of Rp-ds*mlpt* nymphs allows a better understanding of vector-parasite interaction

The parasite *Trypanosoma cruzi* colonises the digestive tract of *R. prolixus*, where it undergoes proliferation and metacyclogenesis [39]. Several factors are essential for metacyclogenesis, including nutritional intake [52] and changes in ecdysone levels [53]. Furthermore, it is well known that neuroendocrine manipulation affects both vector and parasite development, with ecdysone signaling playing a particularly important role in these processes [54,55].

Normally, a nutrient-poor environment such as the hindgut lumen stimulates metacyclogenesis of *T. cruzi* [39,56]. However, we observed a significant increase in the number of epimastigotes and a sharp drop in the number of metacyclic parasites in the hindguts of Rp-*mlpt* silenced nymphs at 21 DAF. We suggest that the phenotypic features of Rp-*mlpt* silenced insects may interfere with *T. cruzi* life cycle in at least two distinct ways: (i) the impairment of midgut peristalsis affects nutrient availability, and consequently interferes with parasite proliferation and differentiation; (ii) the interference with moulting markedly affects the intestinal structure, which, in turn, will have a negative impact on metacyclogenesis.

The Rp-ds*mlpt* nymphs infected by *T. cruzi* presented similar weight profiles and total AM protein content over time after blood feeding with trypanosomes. The exception was the haemozoin profile on the 21st day after infection, where nymphs silenced for Rp-*mlpt* accumulated more haem crystals in the PM. Here we show that Rp-*mlpt* silenced nymphs present an impairment of intestinal muscle fibers, peristalsis and, consequently, on content trafficking. Since proliferation of epimastigotes requires nutritional support [57], a leakage of nutrients from the AM to the PM may occur due to the breakdown of intestinal muscle layers. This extended time of nutrient availability in the PM would allow for a complete digestion of haemoglobin and release of haem. Haem could sustain proliferation of epimastigotes, as it is well-known that *T. cruzi*, like other trypanosomatids, must import haem from their hosts to proliferate [58]. The complete digestion of haemoglobin may also explain the higher haemozoin content in Rp-*mlpt* silenced nymphs when compared with the controls, since haemozoin crystals constitute 97% of all iron species found in the midgut of *R. prolixus* [59,60]. The complete digestion of haemoglobin in Rp-ds*mlpt* nymphs would also lead to the lack of important haemoglobin-derived fragments. For instance, Garcia and colleagues [52] demonstrated that fragments derived from haemoglobin degradation by proteolytic activity in the PM are important modulators of *T. cruzi* metacyclogenesis. Lastly, the accumulation of unprocessed primary urine may act as an additional nutritional support for the proliferation of *T. cruzi* epimastigotes [61], since amino acids might be delivered into the hindgut.

The scarcity of metacyclic trypomastigotes in Rp-ds*mlpt* nymphs can be at least partially explained by the disruption of the moulting process. It is well known that the rectal cuticle of insects is completely renewed during moulting [62]. Importantly, the attachment of *T. cruzi* epimastigotes to the rectal cuticle of triatomines is an essential step that precedes metacyclogenesis [63]. In this context, Mendonça-Lopes and colleagues [64] have recently demonstrated that the physiological levels of ecdysone are essential to the normal development of the rectal cuticle in *R. prolixus*, including the formation of a luminal hydrophobic wax layer, which is recognized by epimastigotes as the substrate to adhesion before starting the metacyclogenesis. Interestingly, the disruption of ecdysone signalling drastically interferes with the cuticle structure, promoting the loss of the superficial wax layer, which leads to the inhibition of parasite attachment [64]. Here we show that moulting is completely abolished in *R. prolixus* after silencing of Rp*-mlpt*. Since *mlpt* expression has been shown to be regulated by ecdysone in *D. melanogaster* [24,27], it is conceivable that the privation of Rp-*mlpt* encoded peptides will potentially interfere with the normal formation of the hindgut cuticle in *R. prolixus*. Abnormal hindgut cuticle would inhibit *T. cruzi* adhesion and drastically reduce the emergence of infective metacyclic trypomastigotes.

The data presented here show that Rp*-mlpt* gene plays a role in post-embryonic phases of *R. prolixus* development, being consistent with previously reports from fruit flies and silkworms [19,37,65]. Overall, this work demonstrates that the *R. prolixus* smORF-containing gene Rp-*mlpt* is essential for digestive physiology, moulting and *T. cruzi* metacyclogenesis in the digestive tract, thus opening new possibilities in the search for new molecular targets for vector control.

## Acknowledgements

R.N.d.F. is a CNPq fellow (311470/2020-3). R.N.d.F was also funded by FAPERJ (E-26/210.119/2022, E-26/210.708/2021, E-26/221.169/2019, E-26/201.093/2020, E-26/202.605/2019 and E-26/210.264/2018). C.A. was a PhD student of PPG-PRODBIO-Macaé (CAPES scholarship) and B.R. is a PhD student at PPG-PCM-ICB-UFRJ. We thank Isabelle Chagas for her help in editing the parasite micrographies.

## Author Contributions

C.A.O.S., S.S.A., F.B.M., B.d-C.-R., J.R.S., and J.L.N-S.: conceptualization of ideas, conducting research and investigation, writing and original draft preparation; L.R., J.A.F.E. and B.R.: conducting research and investigation; C.L.: funding acquisition; R.N.-d-F.: conceptualization ideas, conducting research and investigation, writing and original draft preparation, funding acquisition.

